# DNA methylation profiles in urothelial bladder cancer tissues and children with schistosomiasis from Eggua, Ogun State, Nigeria

**DOI:** 10.1101/2023.05.18.541398

**Authors:** Cephas A. Akpabio, Rachael P. Ebuh, Oluwaseun E. Fatunla, Henrietta O. Awobode, Chiaka I. Anumudu

## Abstract

Squamous cell carcinoma has been attributed to chronic schistosomiasis and is the predominant type of bladder cancer in schistosomiasis endemic areas. The aim of this study was to assess early promoter DNA methylation in selected genes implicated in schistosomiasis-associated bladder cancer (SABC). A total of 161 urine samples were collected from school aged children in Eggua Community of Ogun State and examined by microscopy for *Schistosoma haematobium* eggs. From this sample, a subset of 34(21.1%) urine samples positive for *S. haematobium* eggs and 22 formalin fixed paraffin-embedded bladder cancer tissues obtained from the University College Hospital Ibadan, were subjected to DNA isolation and bisulfite DNA conversion. Quantitative methylation specific PCR was used to determine the methylation status of *APC, RARβ2, RASSF1A* and *TIMP3* in the samples. Methylation in *APC, RARβ2, RASSF1A and TIMP3* was observed in 24(70.6%), 18(52.9%), 15(44.1%) and 8(23.5%) of the positive urine samples respectively and in 7(31.8%), 13(59.1%), 17(77.3%) and 8(36.4%) of bladder cancer tissues respectively. *APC, RARβ2* and *RASSF1A* were 5-fold, 2-fold and 27-fold downregulated respectively in positive urine samples and 9-fold, 3-fold and 15-fold downregulated respectively in the bladder cancer tissues. The odds of promoter methylation in *RARβ2* (OR: 1.133) were likely even with light infection. Gene promoter DNA methylation in tumour suppressor genes was observed in schistosomiasis cases. Hence, DNA methylation may occur during active *Schistosoma haematobium* in children. This result may serve as an early non-invasive biomarker to detect and hint at the risk of developing SABC later in life.

**Author summary:** *Schistosoma haematobium* can survive in the host for more than 20 years, during which time it causes damage to the bladder tissues and sometimes with no symptoms. Immune response to the parasite infection is inflammatory and leads to several morbidities like anaemia, undernutrition, dysuria, and female genital sores and may result in malignant transformation (schistosomiasis-associated bladder cancer) which presents in later years. Children are more susceptible to schistosomiasis because of having a naive immune system, making them targets for these morbidities, and including the possibility of developing bladder cancer in later years. DNA methylation which is often the first step in malignant transformation is known to be induced by inflammation during chronic schistosomiasis. Hence, assessing DNA methylation can serve as a biomarker for predicting the risk of developing bladder cancer later in life. In this study, we have established that DNA methylation occurs during childhood schistosomiasis and represents the time when events leading up to malignant transformation may begin. We suggest that once there is a schistosomiasis infection, DNA methylation will occur and unless the disease is treated on time, the individual is at risk of malignant transformation later in life.

## Introduction

### Schistosomiasis

Schistosomiasis is a neglected tropical disease (NTD) that is endemic in sub-Saharan Africa, Middle East and Asia [1]. It is a water-borne disease caused by species of the genus *Schistosoma*. The species that are endemic to Africa are *S. haematobium* and *S. mansoni* which cause urogenital schistosomiasis and intestinal schistosomiasis respectively. Individuals get infected through contact with water bodies that have been contaminated with infective larval stage (cercaria) of the parasite. Data from the World Health Organization [2], show that schistosomiasis is present in 78 countries, and it is estimated that at least 90% of people requiring treatment for schistosomiasis reside in Africa. It has been estimated that over 700 million people worldwide in endemic regions are at risk of infection, with over 200 000 deaths annually [3].

The host immune response to the various stages of the life cycle of *Schistosoma haematobium* is inflammatory in nature. The consequence of the inflammatory response is granulomatous formation around eggs lodged in the tissues of the bladder. The granulomata usually conjugate, forming tubercles and nodules that often ulcerate [4]. In chronic schistosomiasis, this leads to several morbidities such as anaemia, undernutrition, dysuria and female genital sores [5]; and may result in malignant transformation (squamous cell carcinoma) which usually presents at a late stage [6].

In addition, lesions due to schistosomiasis are assumed to play a part in intensifying the exposure of the bladder epithelium to mutagenic substrates from tobacco or other known toxic chemicals. Although the mechanisms underlying the link between schistosomiasis and bladder cancer are still debatable, available data shows that schistosomiasis induces DNA methylation via chronic inflammation. DNA methylation is known to be involved in numerous cancer phenotypes.

### Schistosomiasis-induced DNA Methylation and Bladder Cancer

DNA methylation is the most studied epigenetic modification and is the process in which a methyl group forms a covalent bond with 5′ position of a cytosine ring forming 5- methylcytosine (5mC). This event usually occurs in CG rich regions called CpG islands. These are regions that have more cytosines followed by guanines than other nucleotides. CpG islands are upstream, and make the bulk of gene promoter regions, in addition to playing crucial roles in gene expression. Methylated promoter regions recruit other methylation enzymes, thereby making access by transcription factors more difficult. Naturally, the process of DNA methylation is carried out by a group of DNA methyltransferase enzymes (DNMTs) namely; DNMT3a and DNMT3b and DNMT1 [7]. The former group is responsible for *de novo* DNA methylation during embryonic development, while DNMT1 maintains the methylation in subsequent cell division using hemimethylated strands [8]. This allows for normal gene regulation and expression.

In contrast, DNA methylation patterns can be altered directly or indirectly by disease causing pathogens (bacteria, parasites and viruses) [9] or chemical agents like amines, tobacco and arsenic. This has debilitating effects on gene expression and function; thus, impacting disease progression including cancer and resistance to therapy. Aberrant DNA methylation (hyper-and hypomethylation) has been associated with numerous diseases including cancer. Many cancerous cells are characterized by different abnormal DNA methylation patterns, which are usually distinct and can be used to discriminate between cancerous and normal cells.

Chronic schistosomiasis which synchronizes with chronic inflammation occurs during active infection in children and peaks at late adolescence [10]. It has also been established that chronic inflammation elicits aberrant DNA methylation, and that the latter is believed to constitute the initiation phase in cancer development. Based on these observations, it can be suggested that early events (DNA methylation) leading up to SABC may occur in children with active schistosomiasis infection. Therefore, assessing DNA methylation patterns in infected children may aid in the identification of individuals who may be at risk of developing SABC later in life, and provide the basis for early intervention.

Schistosomiasis Associated Bladder Cancer (SABC) has not been documented in Eggua, southwest Nigeria, but Onile *et al*. [11] reported that about 62% of bladder pathologies, detected by ultrasonography were associated with chronic schistosomiasis, suggesting altered DNA methylation patterns would have occurred. In addition there are reports of cancer-specific DNA methylation alterations in pre-diagnostic blood collected more than 10 years before diagnosis of chronic lymphocytic leukemia [12]. Based on these reports and the fact that children are more susceptible to *Schistosoma haematobium* infection, it can be suggested that epigenetic changes begin early in children during active infections. Thus, they are hotspots for events leading up to disease progression, bladder pathologies and risk of developing schistosomiasis associated bladder cancer later in life.

### DNA Methylation as biomarker for SABC

The gold standard for treatment of bladder cancer is cytoscopy. It is invasive, painful and less effective for SABC. DNA methylation biomarkers have been widely used for early diagnosis, prognosis and prediction of diseases including cancer. These biomarkers are easily (not invasive and not painful) obtainable from body fluids. Different DNA methylation biomarkers have been assessed for detection, prognosis and treatment of SABC which usually present late [6]. It is known that the best form of treatment is prevention, and that DNA methylation events (which are reversible) can occur years before diagnosis of neoplasm, hence it is pertinent to assess these events at its earliest onset. Therefore, the use of non-invasive DNA methylation biomarkers to identify early events in childhood schistosomiasis, which precede SABC later in life, may help speed up intervention measures and treatments to children with chronic infections and at risk of malignant transformation.

This is the first attempt to evaluate DNA methylation induced my *S. haematobium* infection in children. We also look to establish a possible link between *S. haematobium* infection especially in children and SABC. This study will contribute additional knowledge to the understanding of epigenetic changes occurring in schistosomiasis that may predispose the infected individuals, especially during childhood to the risk of developing SABC. Evaluation of the methylation status will help to determine which urine biomarkers can be used, and are effective in hinting at the risk of developing SABC.

This study is aimed at evaluating promoter DNA methylation, as an early non-invasive potential biomarker in children infected with *Schistosoma haematobium* in Eggua Community, and how it can be used to hint on, and identify individuals at risk of developing schistosomiasis-associated bladder cancer later in life. We have been able to establish that during childhood schistosomiasis, DNA methylation is induced especially in tumour suppressor genes. Therefore, once there is schistosomiasis infection and the disease is left untreated, the individual is at risk of developing schistosomiasis-associated bladder cancer later in life.

## Materials and Methods

### Study design

For this study, a cross sectional design was used to collect urine samples once from voluntary participants who were all school aged children. A total of 161 participants between the ages of 5 and 16 years were recruited from Eggua Community, using informed consent procedures. A total of 22 Formalin fixed paraffin-embedded bladder cancer tissue blocks (SABC and Non-SABC) were collected from the cancer registry of the University College Hospital (UCH), University of Ibadan. The cancer tissue blocks were analysed by a Histopathologist from the University College Hospital (UCH), University of Ibadan, to ascertain they were bladder cancer tissues and histological type.

### Ethical Approval

Ethical clearance was obtained from the UI/UCH Health Research Ethics Committee, College of Medicine, University of Ibadan (IRB number: UI/UCH/22/0036). Using informed consent procedures, the assent and consent of the children and their parents/guardians respectively were obtained in writing before sampling was carried out.

### Study Area

Eggua is a rural agricultural community (07°01.592 N; 002°55.083 E) in Yewa North Local Government Area, Ogun State Nigeria. The main sources of water in the area are flowing Rivers, especially River Yewa, which are used for domestic purposes, including drinking, washing and cooking in addition to fishing and swimming. Schistosomiasis is known to be prevalent in this community [13] and bladder pathologies associated with Schistosomiasis has been reported as well [11].

### Study Population

The study population was drawn from school aged children living in the study area. Urine samples from participants positive for schistosoma eggs and the cancer tissue blocks were used as cases while samples from participants without haematuria or schistosoma eggs detected in urine samples were used as controls.

### Sample Size Determination

This was determined according to Charan and Biswas, [14]; and calculated using the formula;

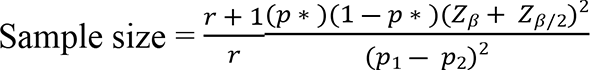

Values for proportions were obtained based on the number of cases and control from a previous study, [15]. Thus number of cases = 57, number of control = 13, r = 0.23, expected proportion in cases (p1) = 0.81, expected proportion in control (p2) = 0.19, Z_β_ = 1.28, Z_β/2_ = 1.96, p* = 0.5.

Therefore, samples from at least 37 participants were used in this study.

### Biological Sample collection and analysis

20ml of voided urine were collected from each participant in reagent bottles. Urinalysis reagent strips (Rapid Labs, UK) were used for rapid detection of blood and analytes, including glucose, ketone, specific gravity, pH, proteinuria and leukocytes in the urine samples, following the manufacturer’s instructions. Samples were immediately centrifuged at 3000 *g* for 10 minutes and the sediments were carefully examined by microscopy for presence of *Schistosoma* eggs. Urine samples were then stored at − 30°C until DNA extraction was carried out. Tissue blocks of formalin fixed paraffin-embedded specimens of Schistosoma-associated bladder cancer and Urothelial bladder cancer were obtained from the bladder cancer registry of UCH, University of Ibadan. Genomic DNA was extracted from them.

### Gene Selection for Promoter Methylation

Four genes (*APC, RARβ2, RASSF1A* and *TIMP3*) associated with schistosomiasis induced bladder pathology and cancers were selected from literature for analysis of methylation abnormalities. These genes have been shown to be highly sensitive as biomarkers for bladder cancer. *RASSF1A, APC* and *RARβ2* are all tumour suppressor genes while *TIMP3* plays a key role in activating apoptotic cascade.

### Primer Design for APC, RARβ2, RASSF1A and TIMP3

The gene ID and organism of the target genes were obtained from the National Centre for Biotechnology Information (NCBI) database. This was then used to search for the promoter region of the gene in the Eukaryotic Promoter Database (EPD). The sequence of the promoter region for each target gene was then retrieved from the database using the sequence retrieval tool. A blast run for the promoter sequence was carried out using the NCBI blast tool in order to determine specificity of the promoter sequence. Primer design specific for methylation assay was then carried out using MethPrimer as previously reported [16].

### Genomic DNA Extraction from bladder cancer tissues and urine samples

For genomic DNA extraction, four tissue sections (≤ 20 µm thick) were obtained from each bladder cancer tissue block, using a microtome and transferred to 1.5 ml microcentrifuge tubes for deparaffinization.

### Deparaffinization of tissue sections and DNA extraction

This was carried out using the Zymo Research Quick-DNA FFPE kit, following the manufacturer’s instructions. Briefly, 400μL of Deparaffinization solution was added to 1.5ml microcentrifuge tubes, incubated at 55°C for 1 minute and thereafter vortexed briefly. Deparaffinazation solution was then discarded. Deparaffinized tissue digested using Proteinase K, dH_2_O and 2X digestion buffer. This was incubated at 55°C overnight for 12-16 hours. DNA was purified and eluted with 70μL of DNA Elution Buffer. Purified DNA was then stored at ≤-20°C for further use.

### DNA Isolation from urine samples

34 urine samples positive for Schistosoma eggs were used for DNA extraction. DNA was also extracted from 16 urine samples that were microscopically negative for Schistosoma eggs and this served as control. This was done using the Geneaid gDNA Microkit, following the manufacturer’s instructions. Briefly, 1ml of urine was transferred to a 1.5ml microcentrifuge tube and centrifuged at 6000xg for 2 minutes and the supernatant discarded. This was resuspended using 500μL of elution buffer, centrifuged at 600xg for 1 minute and the supernatant discarded. Cells were lysed with 200μL of S1 buffer and 20μL of Proteinase K and incubated at 60°C for 30 minutes. Further lysis was done with 200μL of S2 buffer and incubated at 60°C for 20 minutes. DNA was bound using 200μL of absolute ethanol, washed and eluted with 60μL of DNA elution buffer. Eluted DNA was stored at ≤-20°C for further use.

### Bisulfite Treatment of Isolated DNA

DNA extracted from urine samples and paraffinized tissue were subjected to bisulfite treatment, which converts unmethylated cytosine residues to uracil residues, leaving the methylated cytosines as previously reported [17]. Briefly, 50μL of genomic DNA from each sample were denatured using 2μl of freshly prepared NaOH (final concentration, 3M) and incubated 37°C for 10 minutes. The denatured DNA was then mixed with freshly prepared sodium bisulfite solution (final concentration, 5M), covered with mineral oil and incubated in the dark at 50°C for 16 hours. The heavy mineral oil was carefully separated from the reaction solution and the bisulfite-modified DNA were purified using GenepHlow Gel/PCR purification kit, following manufacturer’s instructions. Modified DNA was then stored at − 80 °C.

### Real Time Quantitative Methylation Specific PCR (qMSP) for Bisulfite Converted DNA

The qMSP were conducted for each bisulfite-modified DNA on Applied Systems Bio Rad iQ5 Thermocycler. Amplification was carried out in a 25µl total reaction volume containing 4µl of master mix (5x HOT FIREPol EvaGreen qPCR Supermix), 0.4μl each of forward and reverse primers for each gene (*APC, RARβ2, RASSF1A and TIMP3*), 18.2µl of nuclease free water and 2μl of modified DNA. Amplification reaction conditions were as follows; hot start at 95°C for 5 minutes, subsequent denaturation at 95°C for 15 minutes, annealing at 48 - 49°C for 15 minutes, extension at 72°C for 20 minutes final extension step at 55°C for 10 minutes. This was carried out for 40 cycles. Amplification was carried out for methylated and unmethylated primers of each gene separately. Melt curves were also generated for each reaction. No template control (NTC) was used as negative control. Level of CpG island methylation for promoter region of each gene was obtained among three categories of samples i.e. infection only (urogenital schistosomiasis alone), bladder cancer and controls (no infection/urogenital schistosomiasis). Primer sequences, their parameters and qMSP conditions are shown in supplementary data S1 Table.

### Interpretation and Data Analysis for qMSP

Average melt temperature was used to ascertain actual amplified product for each primer. Relative quantification normalized against unit mass (number of samples) was used to determine fold change with the control sample as the calibrator. This was determined using; Ratio_(test/calibrator)_ = 2^ΔCt^. Where ΔCt = Ct(calibrator) – Ct(test).

## Statistical Analysis

All statistical analysis was done using SPSS version 20. Pearson’s Chi Square test was used to test association between promoter region methylation of each gene with age, gender and cancer histological type. Odds ratio was used to predict the risk of promoter methylation in relation to the intensity of infection. Receiver Operating Curve (ROC) analysis was used to assess promoter region hypermethylation of the selected genes to distinguish between cases and controls.

## Results

A total of 161 school aged children participated in this study. Out of this, 78(48.5%) were males while 83(51.5%) were females, with a mean age of 10.7 years (Table 1). A total of 34 urine samples were positive for *Schistosoma haematobium* (*Sh*) eggs after microscopic examination for a prevalence of 21.1%. Of this, light infection (<50 eggs/10mL of urine) occurred in 32(94.1%) samples, while heavy infection (≥50 eggs/10mL of urine) was seen in 2(5.9%) samples. Urinalysis showed that 33(20.5%) of the participants had haematuria, out of which 11(33.3%) were positive for *S. haematobium* infection (Table 1). Equal number (17) of males (21.8%) and females (20.5%) had *S. haematobium* infection (Table 1).

**Table 1:** Demographic Characteristics of Participants and Bladder Cancer Tissues.

| Parameters | Number |  | Total (%) |
| --- | --- | --- | --- |
|  | Infected (%) | Uninfected(%) |  |
| Participants |  |  |  |
| Sex |  |  |  |
| Male | 17(21.8) | 61(78.2) | 78(48.5) |
| Female | 17(20.5) | 66(79.5) | 83(51.5) |
| Total | 34(21.1) | 127(78.9) | 161(100) |
| Age Group |  |  |  |
| 5 – 10 | 13(22.4) | 45(77.6) | 58(36.0) |
| 11 – 16 | 21(20.8) | 80(79.2) | 101(62.7) |
| Missing data | 0(0) | 2(100) | 2(1.2) |
| Total | 34(21.1) | 125(77.6) | 161(98.7) |
| Haematuria |  |  |  |
| Yes | 11(33.3) | 22(66.7) | 33(20.5) |
| No | 23(18.0) | 105(82.0) | 128(79.5) |
| Total | 34(21.1) | 127(78.9) | 161(100) |
| Bladder Cancer Tissues |  |  |  |
| Sex |  |  |  |
| Male |  |  | 12(54.6) |
| Female |  |  | 5(22.7) |
| Missing data |  |  | 5(22.7) |
| Total |  |  | 22(100) |
| Age Group |  |  |  |
| <55 |  |  | 7(31.8) |
| ≥55 |  |  | 9(40.9) |
| Missing data |  |  | 6(27.3) |
| Total |  |  | 22(100) |
| Tumor Type |  |  |  |
| UC |  |  | 13(59.1) |
| Invasive UC |  |  | 3(13.6) |
| Papillary UC |  |  | 1(4.6) |
| Invasive Papillary UC |  |  | 2(9.1) |
| Invasive SCC |  |  | 2(9.1) |
| Invasive Adenocarcinoma |  |  | 1(4.6) |
| Total |  |  | 22(100) |

A total of 50 urine samples were then subjected to further analysis of DNA methylation; 34 were positive for *Schistosoma haematobium* (*Sh*) eggs and 16 negative for both *S. haematobium* eggs and haematuria and thus, served as controls.

Archived FFPE bladder cancer tissues composed 12(54.6%) males while females were 5(22.7%) with an average age of 55.8 years. More males (54.6%) than females (22.7%) had bladder cancer (Table 1). Urothelial carcinoma was the most common type of tumour, constituting 59.1% of the samples (Table 1).

### Gene Promoter Methylation of the Target Genes in Schistosomiasis and Bladder Cancer

Gene promoter methylation of the target genes was assayed using a total 56 samples (34 *Sh* positive urine and 22 bladder cancer tissues). Promoter methylation in the following genes, *APC, RARβ2, RASSF1A and TIMP3* were evaluated. *RASSF1A* was methylated in more (32(57.1%) of all the samples, while *TIMP3* was the least methylated in 16(28.6%) of samples (Table 2). Figure 1 shows the images of the qPCR amplification cycle and melting temperatures for *APC* gene. The images of the qPCR amplification cycle and melting temperatures for *RARβ2, RASSF1A and TIMP3* are shown in appendix figures I, II, and III respectively.

**Fig 1:**
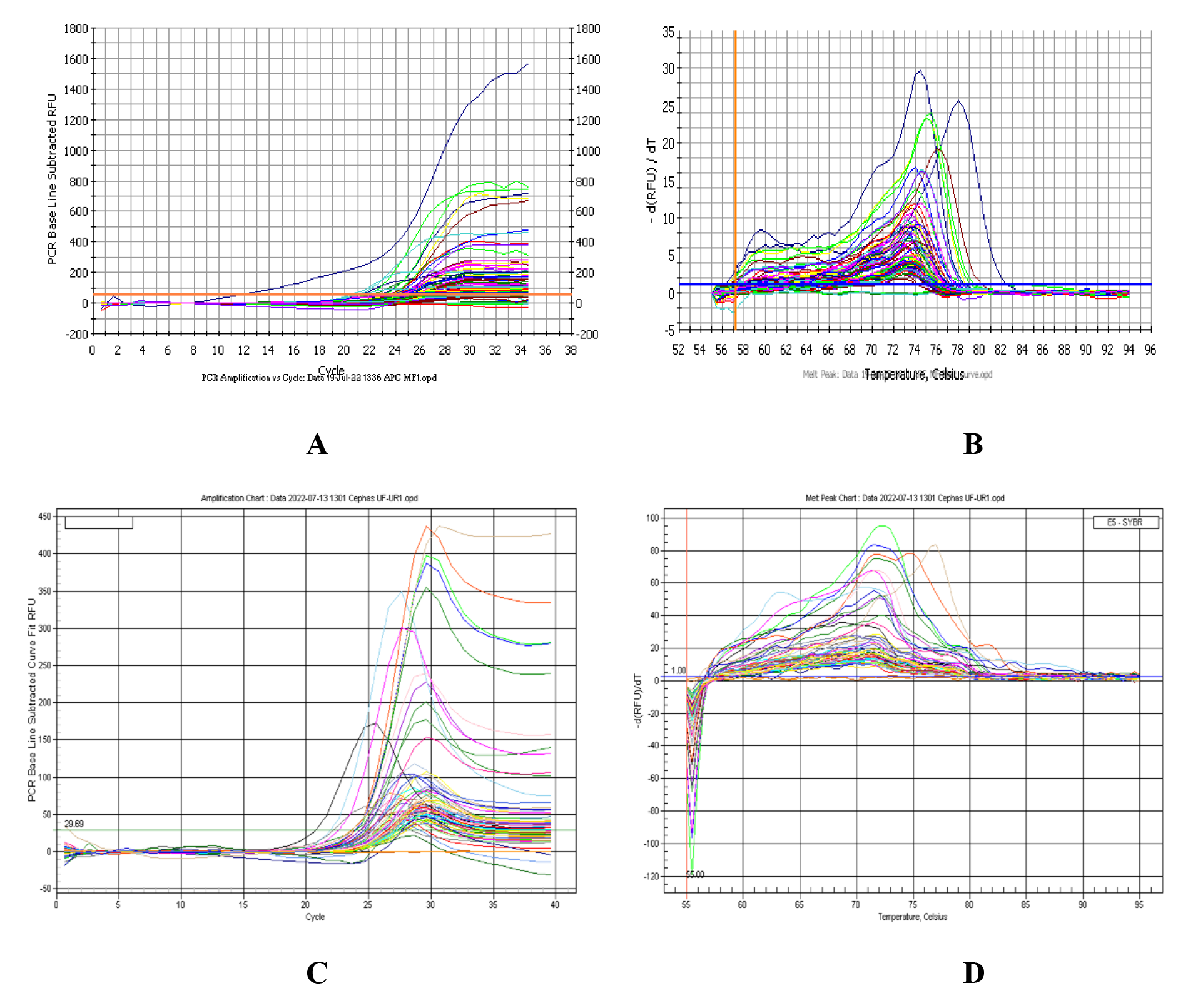
qPCR amplification and melting temperatures for *APC*: A) qPCR amplification cycle for methylation reaction; B) Melting temperature for methylation reaction; C) qPCR amplification cycle for unmethylation reaction; D) Melting temperature for unmethylation reaction

**Table 2:** Promoter methylation of *APC*, *RARβ2*, *RASSF1A* and *TIMP3* genes in schistosomiasis and cancer tissues

| GENE | SAMPLE |  | Total<br>n(%) |
| --- | --- | --- | --- |
|  | Cancer Tissue<br>n(%) | *Schistosoma haematobium*<br>Positive Urine<br>n(%) |  |
| *APC* |  |  |  |
| Methylated | 7(31.8) | 24(70.6) | 31(55.4) |
| Unmethylated | 15(68.1) | 10(29.4) | 25(44.6) |
| *RARβ2* |  |  |  |
| Methylated | 13(59.1) | 18(52.9) | 31(55.4) |
| Unmethylated | 9(40.9) | 16(47.1) | 25(44.6) |
| *RASSF1A* |  |  |  |
| Methylated | 17(77.3) | 15(44.1) | 32(57.1) |
| Unmethylated | 5(22.7) | 19(55.9) | 24(42.9) |
| *TIMP3* |  |  |  |
| Methylated | 8(36.4) | 8(23.5) | 16(28.6) |
| Unmethylated | 14(63.6) | 26(76.5) | 40(71.4) |

In comparing both bladder cancer tissues and positive urine samples, there were more promoter methylation in *APC* (24) and *RARβ2* (18) in UGS samples than cancer tissues: *APC* (7) and *RARβ2* (13); whereas promoter methylation was lower for *RASSF1A* (15) in UGS than for *RASSF1A* (17) in cancer tissue. The number of samples with promoter methylation for *TIMP3* was the same (8), for both UGS and cancer tissues (Table 2).

In the positive urine samples, *APC, RARβ2* and *RASSF1A* were 5-fold, 2-fold and 27-fold hypermethylated respectively, whereas *TIMP3* had no significant fold change. When compared to the cancer tissues, APC and *RARβ2* were more hypermethylated with a 9-fold and 3-fold change respectively, *RASSF1A* showed lesser hypermethylation with a 15-fold change, while *TIMP3* equally had no significant fold change (Figure 2).

**Fig 2:**
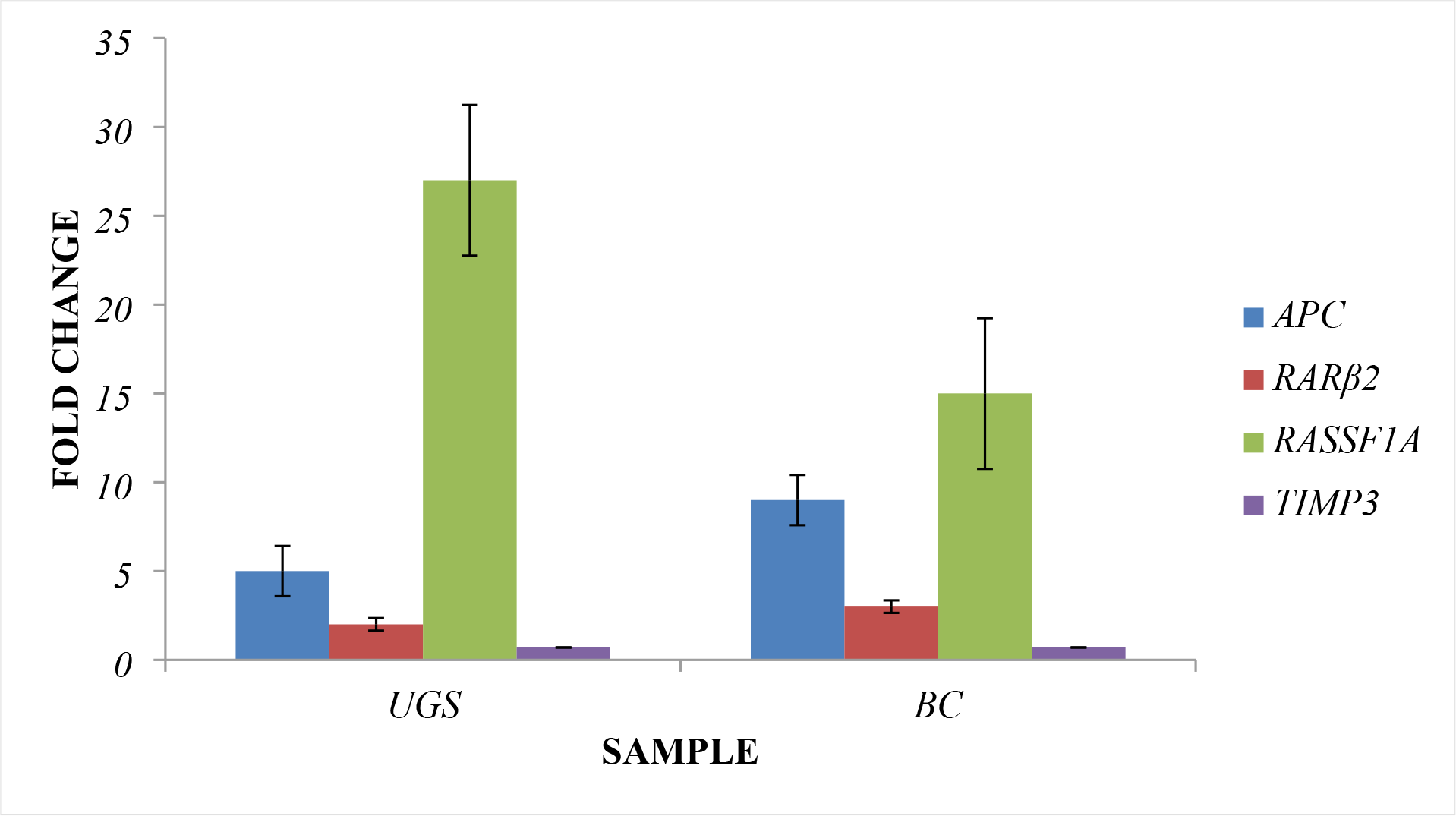
Fold change in promoter methylation of tested genes was highest in *APC and RASSF1A* **Legend: UGS = urogenital schistosomiasis, BC = bladder cancer**

Pearson’s Chi-square test was used to test the association between the promoter methylation of the genes of interest and age group, gender, intensity of infection for schistosomiasis (Table 3) and age group, gender and histological type for bladder cancer (Table 4).

**Table 3:** Association of promoter methylation of *APC*, *RARβ2*, *RASSF1A* and *TIMP3* with urogenital schistosomiasis

| Gene | Parameters | Number Infected | Methylation(%) | *p*-value |
| --- | --- | --- | --- | --- |
| *APC* | **Age Group** |  |  | 0.555 |
|  | 5 – 10 | 13 | 9(69.2) |  |
|  | 11 – 16 | 21 | 15(71.4) |  |
|  | **Total** | **34** | **24(70.6)** |  |
|  | **Sex** |  |  | 0.586 |
|  | Male | 17 | 12(70.6) |  |
|  | Female | 17 | 12(70.6) |  |
|  | **Total** | **34** | **24(70.6)** |  |
|  | **Intensity of Infection** |  |  | **0.000\*** |
|  | Light | 32 | 22(68.8) |  |
|  | Heavy | 2 | 2(100) |  |
|  | **Total** | **34** | **24(70.6)** |  |
| *RARβ2* | **Age Group** |  |  | 0.449 |
|  | 6 – 10 | 13 | 6(46.2) |  |
|  | 11 – 16 | 21 | 12(57.1) |  |
|  | **Total** | **34** | **18(52.9)** |  |
|  | **Sex** |  |  | 0.309 |
|  | Male | 17 | 8(47.1) |  |
|  | Female | 17 | 10(58.8) |  |
|  | **Total** | **34** | **18(52.9)** |  |
|  | **Intensity of Infection** |  |  | **0.001\*** |
|  | Light | 32 | 17(53.1) |  |
|  | Heavy | 2 | 1(50.0) |  |
|  | **Total** | **34** | **18(52.9)** |  |
| *RASSF1A* | **Age Group** |  |  | 0.495 |
|  | 6 – 10 | 13 | 5(38.5) |  |
|  | 11 – 16 | 21 | 10(47.6) |  |
|  | **Total** | **34** | **15(44.1)** |  |
|  | **Sex** |  |  | 0.542 |
|  | Male | 17 | 7(41.2) |  |
|  | Female | 17 | 8(47.1) |  |
|  | **Total** | **34** | **15(44.1)** |  |
|  | **Intensity of Infection** |  |  | **0.010\*** |
|  | Light | 32 | 15(48.9) |  |
|  | Heavy | 2 | 0 |  |
|  | **Total** | **34** | **15(44.1)** |  |
| *TIMP3* | **Age Group** |  |  | 0.256 |
|  | 6 – 10 | 13 | 2(15.4) |  |
|  | 11 – 16 | 21 | 6(28.6) |  |
|  | **Total** | **34** | **8(23.5)** |  |
|  | **Sex** |  |  | 0.256 |
|  | Male | 17 | 6(35.3) |  |
|  | Female | 17 | 2(11.8) |  |
|  | **Total** | **34** | **8(23.5)** |  |
|  | **Intensity of Infection** |  |  | 0.127 |
|  | Light | 32 | 8(25.0) |  |
|  | Heavy | 2 | 0 |  |
|  | **Total** | **34** | **8(23.5)** |  |

\*Significant at $p \leq 0.05$

**Table 4:**
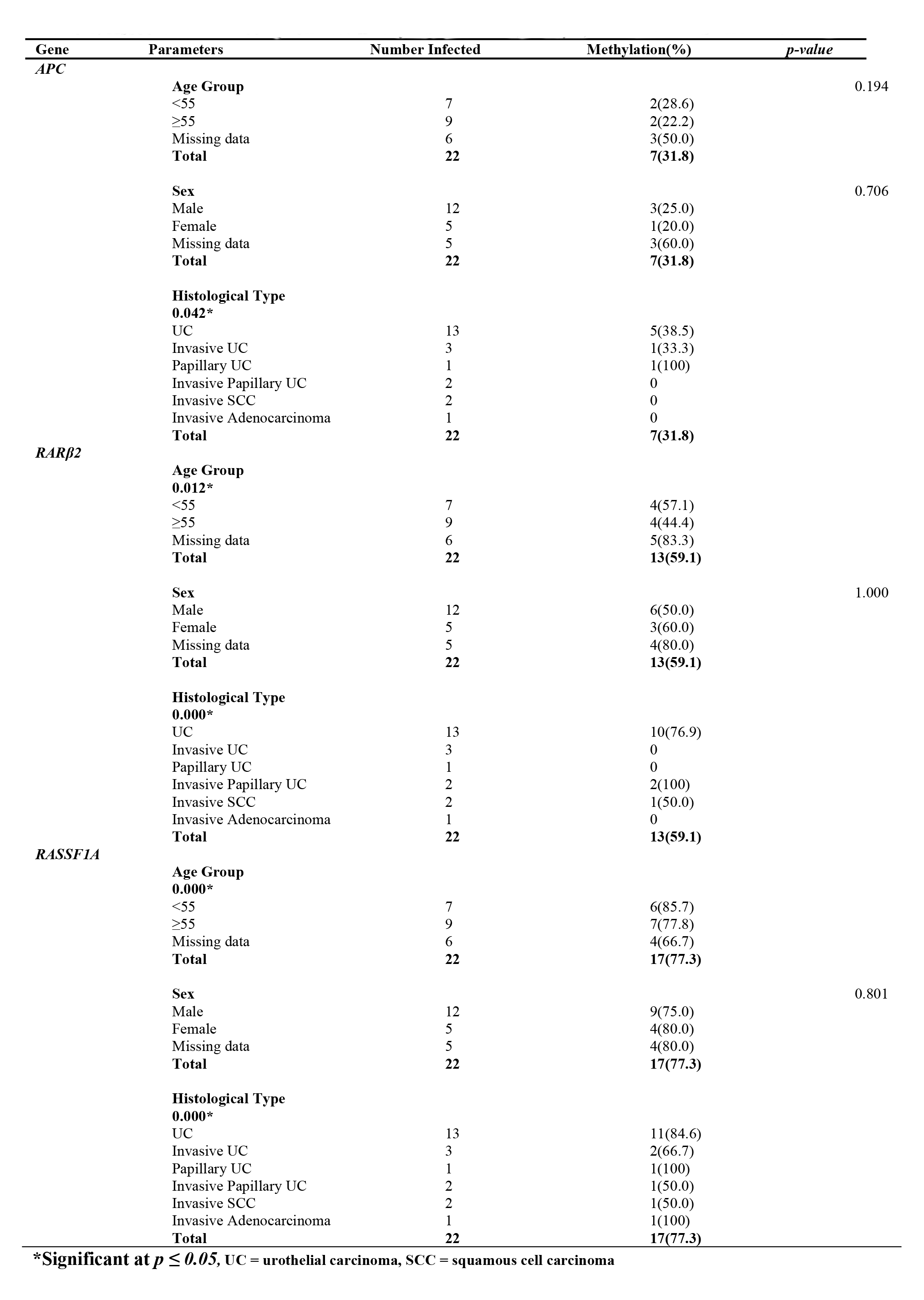
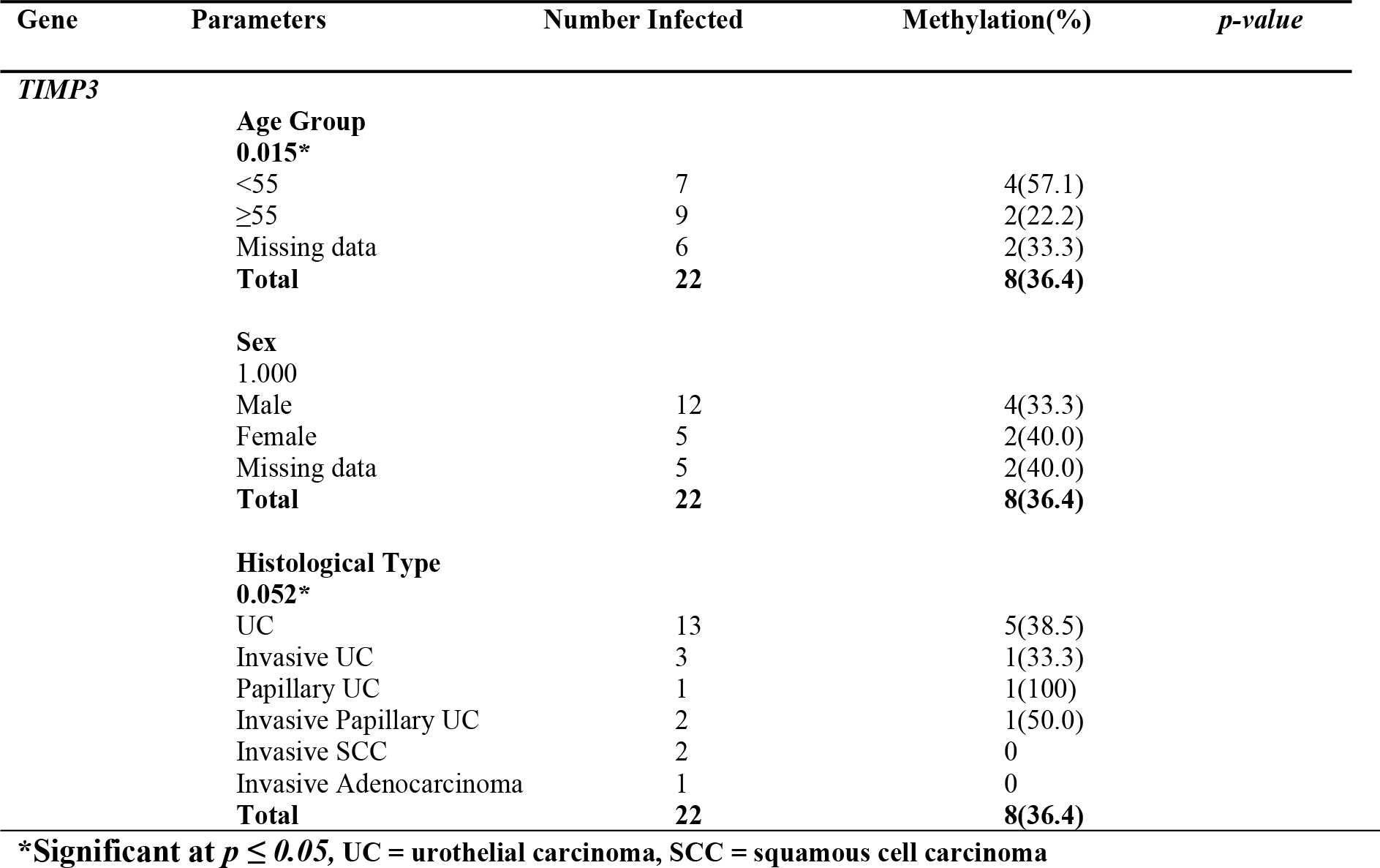
Association between promoter methylation of *APC*, *RARβ2*, *RASSF1A* and *TIMP3* and BC

As shown in Table 3 and Figure 3, majority of the participants with urogenital schistosomiasis belong to the age group 11 – 16 years and this group also accounted for the highest number of promoter methylation of the tested genes, but there was no association between age and promoter methylation (p>0.05). Although majority of bladder cancer patients belonged to the age group ≥55 years, but the age group <55 had highest frequency of promoter methylation of the tested genes. There was a significant association between age of bladder cancer patients and promoter methylation of tested genes except *APC* at p<0.05 (Table 4).

**Fig 3:**
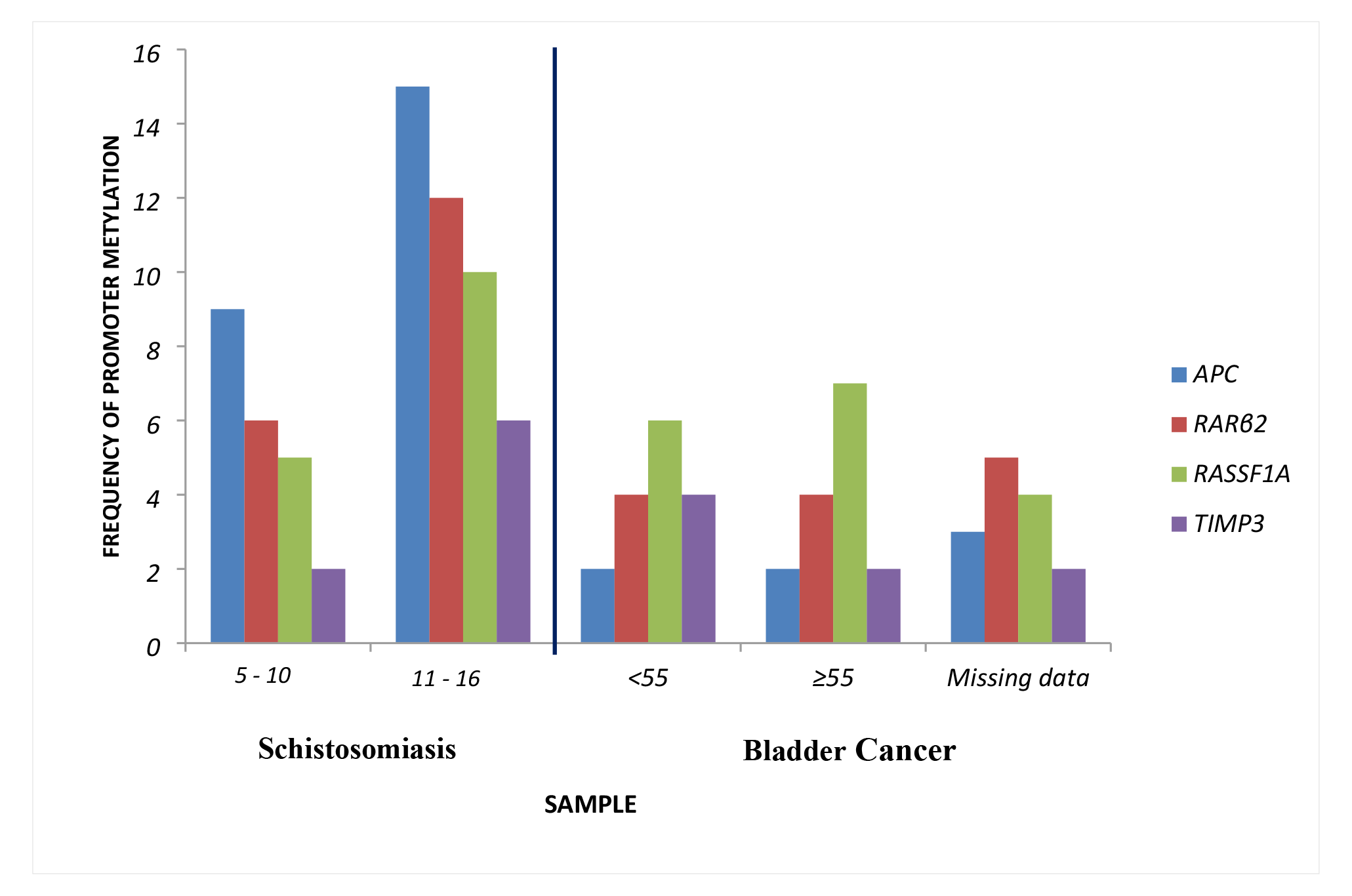
No association between promoter hypermethylation of *APC*, *RARβ2*, *RASSF1A* and *TIMP3* and age

As shown in Table 1, equal numbers of male and female children were infected with *Schistosoma haematobium*. While females had the highest promoter hypermethylation for *RARβ2* and *RASSF1A*, males had highest promoter hypermethylation in *TIMP3* and both had equal number of promoter hypermethylation for *APC* (Table 3). There was no association between sex and promoter methylation in children with schistosomiasis (p>0.05). In the case of bladder cancer tissues used, more males than females had bladder cancer and males also had the highest number of promoter methylation for all tested genes, but there was association between sex and promoter methylation p>0.05 (Table 4).

Intensity of infection was associated with promoter methylation for only *APC*, *RARβ2* and *RASSF1A* at p<0.05 (Table 3). For bladder cancer, histological type was also associated with promoter methylation for all tested genes at p≤0.05 (Table 4).

Odds ratio was used to predict if intensity of infection was associated with gene promoter hypermethylation. Our results showed that promoter hypermethylation was more likely even at light infection for *RARβ2* (Table 5). Receiver Operating Characteristic (ROC) curve was generated to assess the ability of the assay to predict promoter methylation. The ROC analysis for *APC* had sensitivity, specificity and area under curve (AUC) of 55.4%, 75% and 0.908 respectively (Figure 4). The ROC analysis for *RARβ2, RASSF1A, and TIMP3* are shown in appendix figures IV, V, and VI respectively. The ROC analysis for all tested genes had a p-value of p < 0.001.

**Fig 4:**
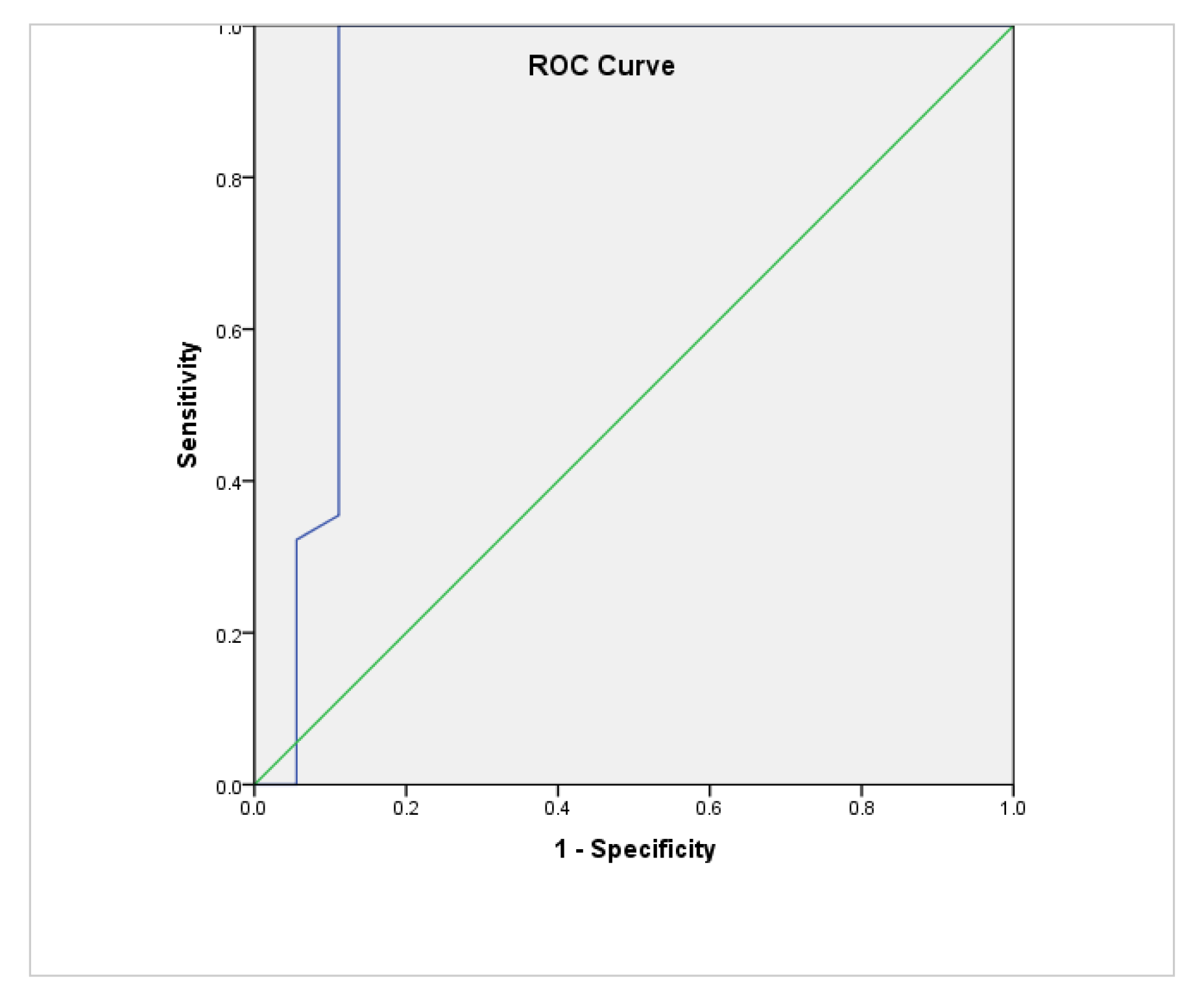
AUC-ROC Curve for assessing promoter methylation of *APC* (sensitivity = 55.4%, specificity = 75%, p<0.001, AUC = 0.908)

**Table 5.** Association between intensity of infection and promoter methylation of *APC*, *RARβ2*, *RASSF1A* and *TIMP3*

| Parameter | Gene | OR | 95%CI |
| --- | --- | --- | --- |
| Intensity of infection<br>(Light/Heavy) | *APC* | .688 | .544 - .868 |
|  | *RARβ2* | 1.133 | .065 – 19.74 |
|  | *RASSF1A* | .531 | .384 - .736 |
|  | *TIMP3* | .750 | .614 - .916 |

## Discussion

Altered DNA methylation patterns can inform, (and have been used) in the diagnosis, prognosis, treatment and management of different cancers and other diseases and disorders [18]. *Schistosoma haematobium* the causative agent of schistosomiasis, a neglected tropical disease, is a designated Class I carcinogen and implicated in SABC. Although its mechanism of action is still debatable, it is believed that *S. haematobium* induces bladder cancer over time via chronic inflammation. Schistosomiasis associated bladder cancer usually presents late and has poor prognosis.

In this study, we proposed that since DNA methylation is elicited by chronic inflammation which is believed to mark the initiation phase of cancers, assessing DNA methylation patterns during active chronic infection in children may hint at the risk of developing SABC at a later stage in life, and provide the basis for early intervention. In the present study, data are presented which showed hypermethylation in the promoter region of tumour suppressor genes. These may suggest that events leading up to SABC begins during active infection in children. This can be used to distinguish between cases and non-cases, and identify individuals at risk of developing SABC later in life.

### Epidemiology of Schistosomiasis in the Study Area

The overall prevalence of urogenital schistosomiasis in this study was 21.1%. This is lower than 30.8% reported for children in the same area by Oladele *et al*. [19]. Meanwhile, Alexander *et al*. [5] reported a lower prevalence of 14% in children in other areas of Ogun State. The low prevalence of urogenital schistosomiasis recorded in this study may be a result of the ongoing mass drug administration in Ogun State. It is reported that at least once a year since 2017, pupils in the state are treated with praziquantel depending on the availability of the drugs [5].

Opara *et al*. [20] reported equal chances of males and females being infected with the parasite. This corroborates the observations in the present study, with equal number (17) of males (21.8%) and females (20.5%) infected with *S. haematobium*. This is however at variance with other reports that observed high infection in males than females [3] and in females than in males [5]. This has been attributed with the socio-cultural practices of the study areas, as well as how frequent individuals get in contact with cercariae infested water bodies.

Intensity of infection is usually associated with chronicity of disease, clinical manifestations, morbidities, as well as bladder pathologies. Most of those who were schistosomiasis positive had only light infection, as has consistently been seen in this area [11] and other parts of the country [5, 20]. Conversely, reports from other parts of Nigeria [10] and Ghana [21] found that majority of participants had heavy infection (≥50 eggs/10mL urine). Only two participants had heavy infections in this study, which is similar to the report of Alexander *et al*. [5].

### Hypermethylation of Gene Promoter Region in UGS during Childhood and UBC

In the current study, the promoter region of *APC*, *RARβ2*, *RASSF1A* and *TIMP3* were hypermethylated in 31(55.4%), 31(55.4%), 32(57.1%) and 16(28.6%) of all the samples analyzed respectively. Of this, the promoter regions of *APC*, *RARβ2*, *RASSF1A* and *TIMP3* were hypermethylated in 24, 18, 15 and 8 respectively of the schistosomiasis positive samples, whereas they were hypermethylated in 7, 13, 17 and 8 respectively of the bladder cancer tissues. The observations of the current study shows that schistosomiasis in children may induce DNA methylation in the target genes due to chronic inflammation. Egg deposition in tissues usually corresponds with inflammatory response. With the observation of eggs in urine of infected children in this study, it can be suggested that chronic inflammation was taking place and may be the reason for the hypermethylation of the target genes.

To ascertain that the target genes are markers for bladder cancer, we tested for promoter region hypermethylation of the target genes in bladder cancer tissues. Our observations showed marked promoter hypermethylation in the target genes, and supports our observation of the same promoter hypermethylation in childhood schistosomiasis. This implies that these genes are markers for bladder cancer. The reports that cancer-specific DNA methylation can occur more than ten years before diagnosis of neoplasm [12] also supports the observation of

DNA methylation in children with schistosomiasis in the current study, and may imply that events leading up to SABC at a later stage in life may begin years before diagnosis. This is in addition to the fact that altered DNA methylation pattern is associated with the initiation phase of cancer or precedes tumorigenesis [22].

Furthermore, there was a significant association between the histological type of the bladder cancer tissues and promoter DNA methylation in *APC* (p<0.005), *RARβ2* (p<0.001), *RASSF1A* (p<0.05) and *TIMP3* (p = 0.05). This implies that DNA methylation of the promoter region of tumour suppressor genes and other genes involved in cell cycle regulation and apoptosis are pre-malignant events that may occur in bladder cancer. This is similar to the report of Zaghloul *et al*. [1], which found DNA methylation of promoter region of these genes and hence suggested their implications for bladder cancer.

### DNA methylation in Childhood Schistosomiasis

In the current study, it was observed that *APC*, *RARβ2* and *RASSF1A* were 5 times, 2 times and 27 times hypermethylated respectively in schistosomiasis positive samples. This was similarly observed in the invasive bladder cancer tissues with a 9-fold, 3-fold and 15-fold hypermethylation for *APC*, *RARβ2* and *RASSF1A* respectively. The presence of DNA methylation in non-tumor urine cells in this study, validated by those in the invasive bladder cancer tissues, may suggest the presence of an “epigenetic field defect”.

Kim and Kim, [22] reports that normal-appearing tissues taken at least 5cm away from invasive tumours had 169 hypermethylated loci; out of which 142 loci were the same as those seen in the invasive tumours. This they described as an “epigenetic field defect”, meaning that methylation was already present before initiation of tumorigenesis. Furthermore, the findings of the present study indicate that *S. haematobium* can effectively induce DNA methylation in non-cancerous exfoliated cells. This is similar to the report of Nakajima *et al*. [23], that *Helicobacter pylori* can effectively induce aberrant DNA methylation in non-cancerous gastric mucosae.

The concept of “field of cancerization” was first reported when it was observed that metachronous primary cancers developed further, even after curative resection in oral cavity cancer [24]. The presence of “epigenetic field defect” may contribute to what is referred to as an “epigenetic field of cancerization”. This implies that aberrant DNA methylation in non-tumour cells may suggest the role of the former in the epigenetic field of cancerization [23]. Therefore, the observations of promoter hypermethylation of tumour suppressor genes in the present study may hint that during schistosomiasis in children, DNA methylation alterations occur, forming an epigenetic field of cancerization which may over time progress to SABC.

The observations of the current study show that the highest number of promoter methylation of the tested genes occurred in children with schistosomiasis in the age group 11-16 years old (Figure 2; Table 3). This may be due to the fact that schistosomiasis infection peaks at adolescence [10], and synchronizes with time when chronic inflammation, restorative hyperplasia, squamous cell metaplasia and DNA lesions may be high [21]. Although there was no association between age and promoter region hypermethylation, these results suggests that DNA methylation events and pre-cancerous lesions may begin during active schistosomiasis infection in childhood.

In this study, there was no association between gender and promoter hypermethylation for tested genes in schistosomiasis positive samples, implying that DNA methylation of promoter region of the tested genes were independent of gender. On the other hand, more males than females had DNA methylation of promoter region of the tested genes in the bladder cancer tissues. This was significant and implied an association between gender and bladder cancer. This may explain why more males than females have bladder cancer [25].

The intensity of infection in this study was significantly associated with promoter DNA hypermethylation of *APC* (p<0.001), *RARβ2* (p = 0.001) and *RASSF1A* (p<0.05). There was no association between intensity of infection and *TIMP3* (p = 0.127). This is similar to a report of association between light intensity and bladder pathologies in adults [11]. Therefore, even in light infections, chronic disease can still occur leading to epigenetic alterations like DNA methylation which may be easily observed than mutations, and precede cytological abnormalities, morbidities, bladder pathologies and the risk of malignant transformation later in life.

Odds ratio was used to predict the risk of gene promoter hypermethylation in relation to the intensity of infection. Our results showed that the gene promoter of *RARβ2* was more likely to be hypermethylated even at light infection, whereas the gene promoter of APC, *RASSF1A* and *TIMP3* were less likely to be methylated at light infection, but in heavy infections. Based on these observations, it can be said that once infection with *S. haematobium* is established, gene promoter methylation will occur. Therefore, it could be suggested that since DNA methylation occurs before or marks the onset of future malignant transformation, childhood schistosomiasis marks the beginning of events leading up to SABC.

The ROC-AUC analysis could be used for routine screening purposes, to ascertain DNA methylation of promoter region of target genes in infected children and to discriminate between cases and non-cases. It could be used to assess DNA methylation fingerprints in individuals who may have had prior infection that have been treated or showing symptoms/morbidity markers of bladder pathologies/cancer. This may help identify individuals at risk of malignant transformation at a later stage in life.

### Possible link between childhood UGS and the risk of developing SABC at a later stage in life

It is known that children under the age of 18 years are the most susceptible group to schistosomiasis infection, and this may be as result of children having a naïve immune system [26]. Infection is highest in children from 5 years old and peaks in adolescents of age between 15 and 18 years [3]. This may explain why there are usually less obvious events that may indicate active infection after adolescence, giving rise to a “concomitant immunity” with a balanced host-parasite inter-relationship that keeps both alive and establishing a reduced parasite’s fertility, limited patient morbidity and resistance to re-infection [4].

Furthermore, it has been established that DNA methylation had occurred in pre-diagnostic blood collected more than 10 years before diagnosis of chronic lymphocytic leukemia [12]. Moreover, the role of epigenetic alterations appears to not only be limited to cancers, and there is even a greater possibility that “epigenetic field defect” may play a part in, and be identified for various diseases [23].

Based on the above observations and the presence of an “epigenetic field defect” in the current study; it can be suggested that DNA methylation (which marks the initiation phase of cancers) begins during active schistosomiasis infection in children under the age of 18 years old. According to Nakajima *et al*. [23], measuring disease risk at a time point using DNA methylation marker will aid individuals change their lifestyles especially for close intensive disease prevention. Therefore, the results of the current study suggest that the epigenetic alterations occurring during childhood schistosomiasis may be the link to SABC at a later stage in life.

These observations imply that early events of DNA methylation of promoter region of tumour suppressor genes (*APC, RARβ2* and *RASSF1A*) and *TIMP3;* involved in apoptosis, observed in this study may occur during active infections of *S. haematobium* in childhood. These alterations if not repaired or incompletely repaired, may be replicated and passed on from one generation of cells to another. This may lead to transformation of cells due to inability of cells to repair damaged cells, as seen in urothelial hyperplasia and squamous cell metaplasia in urine of children with active infection [21]. Urothelial hyperplasia and squamous metaplasia, are themselves potential preneoplastic lesions of the bladder. Subsequently, it may promote the propagation of cells harbouring genotoxic DNA damage and with just a matter of time and further genotoxic damage, before a potential cancer occurs [27].

Thus, hypermethylated genes during childhood infections may be maintained and carried from one generation of cell to another, until infection peaks in late adolescence. At this point, granulomas may have accumulated and gradually replaced by restorative hyperplasia. This may explain why children with UGS have mild bladder pathologies such as thickness of bladder wall and irregularities [28]. These are silent events that are almost not observable and may serve as the link or bridge between chronic UGS in childhood and SABC in later age.

The occurrence of hypermethylation of tumour suppressor genes as seen in the present study is not conclusive that there will be development of cancer. Rather, if disease is left untreated in childhood, there is a higher risk of cancer development in later age. It is evident that epigenetic changes are reversible, therefore DNA methylation induced by schistosomiasis may be reversed if detected early, thus preventing progression or development of disease and/or cancer. Tetteh-Quarcoo *et al*. [29], reports post-praziquantel treatment reversal of cytological abnormalities such as the reversal of transitional metaplastic squamous cells to normal cells after 3-8 weeks in children. Drugs such as DNA methylase inhibitor or histone deacetylase inhibitors are known to restore the activity of suppressed genes.

Therefore, early detection and drugs targeted at reversal of epigenetic changes are areas of research interest. More studies are needed to validate the observations from the present study, especially to check if there is an association between DNA methylation and bladder pathology not just in children with active infection, but adults who may have had schistosomiasis as children or presenting with morbidity markers. Furthermore, this study is limited by the small size. Hence, the need for a large sample size or cohort studies.

This is important because *S. haematobium* may leave behind fingerprints of DNA damage like DNA methylation which may induce carcinogenesis later in life. Some carcinogenic factors are known to leave their fingerprint such as specific DNA methylation patterns in the tissues they damage even if they are no longer present or eradicated, for example *H. pylori*. Moreover, DiNardo *et al*. [30] reports that schistosomiasis-induced CD4+ T cells DNA methylation signature persisted at least 6 months after successful deworming for schistosomiasis. This indicates the continued effect of schistosome infections even after treatment, and is consistent with observations that the pathologies associated with schistosomiasis persists beyond infection [31].

Although it is a schistosomiasis endemic area, it is noteworthy that SABC has not been reported in Eggua community. Although this claim has not been fully verified, there might be few explanations for it. First, as infected individuals grow, they acquire effective immunity to the parasite insult, which may aid in adequate moderation of immune response to the infection. Malignant transformation is usually a result of poorly regulated infection which may lead to fibrosis and SABC. Secondly, during childhood, abnormal cells may have resolved either by themselves or these individuals may have had treatment for the infection. This can be explained by the fact that transitional squamous metaplastic cells are known to resolve with time in children, especially post-praziquantel treatment [29].

Finally, schistosomiasis usually works together with other factors like environmental carcinogens and mutagens to induce bladder cancer [32]. Therefore, these individuals may not be at risk of persistent exposure to these genotoxins at levels high enough to elicit malignant transformation. Nevertheless, further studies are needed to verify these assertions.

## CONCLUSION

The present study has shown that gene promoter region DNA methylation of tumour suppressor genes will occur once *Schistosoma haematobium* infection is established especially in the most vulnerable group (children). The observation of an “epigenetic field of cancerization” in this study could be used as a molecular biomarker to identify individuals at risk of malignant transformation in later age. Based on the presence of an “epigenetic field of cancerization”, it can be suggested that the journey to malignant transformation begins during active infection in childhood, especially if disease is left untreated. Therefore individuals, especially children living in schistosomiasis endemic areas should be given adequate public health attention through adequate diagnosis, mass drug administration, routine follow-up and means of avoiding contact with infested water bodies.

## Acknowledgements

We appreciate the Adele of Ijoun, staff of the primary health care centre in Ijoun and the headteachers of the various schools we obtained samples from.

## Supporting information

S1 Table: Primer sequence and qMSP reaction conditions

S1 Fig. qPCR amplification and melting temperatures for *RARβ2*

S2 Fig. qPCR amplification and melting temperatures for *RASSF1A*

S3 Fig. qPCR amplification and melting temperatures for *TIMP3*

S4 Fig. AUC-ROC Curve for assessing promoter methylation of *RARβ2*

S5 Fig. AUC-ROC Curve for assessing promoter methylation of *RASSF1A*

S6 Fig. AUC-ROC Curve for assessing promoter methylation of *TIMP3*

## Financial disclosure statement

The authors received no specific funding for this work

## Notes

### Competing Interest Statement

The authors have declared no competing interest.

